# Environmentally induced lipidome adaptation in the bacterial model organism *M. extorquens*

**DOI:** 10.1101/650390

**Authors:** Grzegorz Chwastek, Michal A. Surma, Sandra Rizk, Daniel Grosser, Oksana Lavrynenko, Magdalena Rucińska, Helena Jambor, James Sáenz

**Affiliations:** Technische Universität Dresden, B CUBE, Tatzberg 41, Dresden, DE; Lipotype GmbH, Tatzberg 47, Dresden, DE; Łukasiewicz Research Network – PORT Polish Center for Technology Development, Stabłowicka 147, Wrocław, PL; DZD-Paul Langerhans Institute Dresden, Fetscherstraße 74, Dresden, DE; Max Planck Institute of Molecular Cell Biology and Genetics, Pfotenhauerstraße 108, Dresden, DE; Technische Universität Dresden, Medizinische Fakultät, Fetscherstraße 74, Dresden, DE

**Keywords:** Lipidome adaptation, Membrane bioengineering, Lipidome resource, Bacterial-host interactions

## Abstract

Cells, from microbes to man, adapt their membranes in response to the environment to maintain their properties and functions. To adapt, lipid composition is homeostatically regulated to conserve optimal membrane properties. Global patterns of lipidome remodelling are poorly understood, particularly in model organisms with simple lipid compositions that can provide insight into fundamental principles underlying membrane adaptation. Using shotgun lipidomics, we examined the simple yet adaptive lipidome of the plant-associated Gram-negative bacterium *Methylobacterium extorquens* over varying temperature, hyperosmotic and detergent stress, carbon sources, and cell density. We observed that as few as ten lipids account for 90% of the total changes, thus constraining the upper limit of variable lipids required for an adaptive living membrane. Across all conditions, the highest degree of lipidomic variability was observed for changing growth temperature. We also revealed that variations in lipid structural features are not monotonic over a given range of conditions and are heterogeneous across lipid classes. Interestingly, phosphotidylcholine showed the most extreme acyl chain remodeling among all lipid classes, suggesting a new link to its importance in bacterial-host interactions and pathogenicity. These patterns in lipidomic remodeling suggest a highly adaptive mechanism with many degrees of freedom and constrain the lipidomic requirements for an adaptive membrane.

## 1. Introduction

All organisms have at least one membrane that is crucial for compartmentalizing and coordinating biochemical processes within the cell. And yet, nearly a century since the discovery that membranes are made of a lipid bilayer (Gorter and Grendel, 1925) and half a century since integral membrane proteins were proposed in the Fluid Mosaic model (Singer and Nicolson, 1972) we still lack fundamental insight into the design principles required to engineer a functional cell membrane. Much of the membrane’s functionality is associated with the activity of membrane proteins, which perform a diverse range of tasks from signaling to transport. The activity of such proteins is in turn crucially dependent on the biophysical properties of the membrane such as viscosity, thickness, curvature and bilayer asymmetry (Hunte and Richers, 2008; Klose et al., 2013; Pomorski and Menon, 2006; Sanders and Mittendorf, 2011). These properties are largely dictated by membrane lipids (Ernst et al., 2016; van Meer et al., 2008).

Cells produce many lipids with different structures that, in their unique combinations, determine the membrane’s biophysical state. Moreover, cellular lipidomes can vary from tens, for example in the bacterial lipidome reported here, to hundreds of individual lipid species in mammalian cells (Sampaio et al., 2011; van Meer et al., 2008). A major challenge in membrane research is, therefore, to understand how cells regulate individual lipid abundances to maintain an adaptive, functional membrane. Recent progress reconstituting synthetic membranes from the bottom up is paving the way towards the goal of building a self-sustaining adaptive membrane and presently one of the challenges is to implement adaptive lipid remodeling (Brea et al., 2016; Buddingh and van Hest, 2017; Hardy et al., 2015; Tang et al., 2014). Complex lipidomes can be remodeled to modulate collective membrane properties such as viscosity, lipid packing, thickness and phase properties (Klose et al., 2013; Levental et al., 2016). For instance, increasing the proportion of lipids containing double bonds (acyl chain saturation) or changing the number or position of double bonds along the phospholipid acyl chains, can alter the overall membrane viscosity (Fan and Evans, 2015; Ma et al., 2015; Quinn, 1981; Sinensky, 1974).

Singular observations of lipidome composition do not inform which properties the cell membrane senses and adapts to, nor which minimal suite of lipid structure are required for swift adaptation. Lipidomics has provided global insights into the complexity of cellular lipidomes from yeast to man and can provide insight into how cells remodel lipid composition during adaptation (Ejsing et al., 2009). To date, however, there still are only few studies systematically characterizing lipidomic remodeling across a range of conditions and stressors. The most comprehensive study of this kind was performed on yeast (Klose et al., 2012), whose lipidome contains over 150 unique lipids, providing unprecedented insight into the adaptation of a complex eukaryotic lipidome.

Complex organisms possess membranes with diverse and niche-specific functional requirements. While understanding complex lipidomes is important, such systems are not ideal for exploring the fundamental principles underlying lipidome adaptation. Since all membranes require means to homeostatically adapt to perturbations (Kaiser et al., 2011), a simpler organism that is capable of adapting to a broad range of environmental conditions would be a more suitable model system to elucidate the minimal lipidomic requirements for adaptation. Bacteria are an attractive target for this endeavor since some are among the simplest organisms, capable of surviving over a broad range of environments, and many species have been well-characterized and established as model organisms for cell biology.

Here, we characterized the lipidomic adaptivity of the Gram-negative bacterium *Methylobacterium extorquens*. *M. extorquens* has a small lipidome, with 25 phospholipid species as compared with over 100 in *E. coli* (Jeucken et al., 2019). Despite its relatively small lipidome, as a plant- and soil-associated bacterium, *M. extorquens* must adapt to a broad range of chemical and physical conditions (Vorholt, 2012). Furthermore, as a eukaryote-associated bacterium with potential for industrial production of chemicals from methanol, understanding lipidome adaptation in Methylobacterium has broad relevance from bacterial-host interactions to bioengineering. We explored lipidomic remodeling over varying temperature, osmotic and detergent stress, carbon source, and cell density. Surprisingly, we find that most lipidome features remain stable, with only minimal modulations even under significant perturbations. However, other lipidome features are highly plastic and contribute to membrane adaptation. Analysis of structural features such as saturation, acyl chain length or headgroup type revealed signatures of adaptive lipidomes. Globally, we observed that as few as 10 lipids, less than half of all available species, account for the majority of lipidomic adaptation across all perturbations. The patterns of lipidome remodeling that we report here provide a resource for exploring the minimal design principles of living membranes and for designing an adaptive synthetic membrane.

## 2. Results and Discussion

### 2.1 Experimental Design

To determine lipidomic remodeling induced by environmental perturbations, we measured the lipidomes of *M. extorquens* under physical and chemical challenges to cellular membrane homeostasis (Figure 1a; Figure 1-Figure supplement 1). To this end, we varied temperature, osmotic conditions, detergent stresses, carbon sources, and cell densities (Figure 1-Figure supplement 2). Cells were harvested during early exponential growth to minimize their impact on media chemical composition. Temperature was varied from 6 to 30 °C to challenge the viscosity of the membranes. Salt (NaCl) and detergent (Triton X-100) concentrations were varied to produce osmotic and detergent stresses, respectively. Cells were grown on two concentrations of methanol (0.1 and 1% v/v) as the sole carbon source to evaluate the effect of metabolism on lipidomic remodeling. Finally, to characterize lipidomic adaptation at varying cell densities, cells were harvested at early, mid and late exponential, as well as stationary growth stages (Figure 1b). Cellular lipid content was analyzed by high resolution shotgun mass spectrometry (Ejsing et al., 2009) to determine absolute abundances of 25 phospholipids and two hopanoid species, distributed between 5 lipid classes and representing over 90 mol% of the lipidome of *M. extorquens* (Sáenz et al., 2015)(Figure 2).

**Figure 1.**
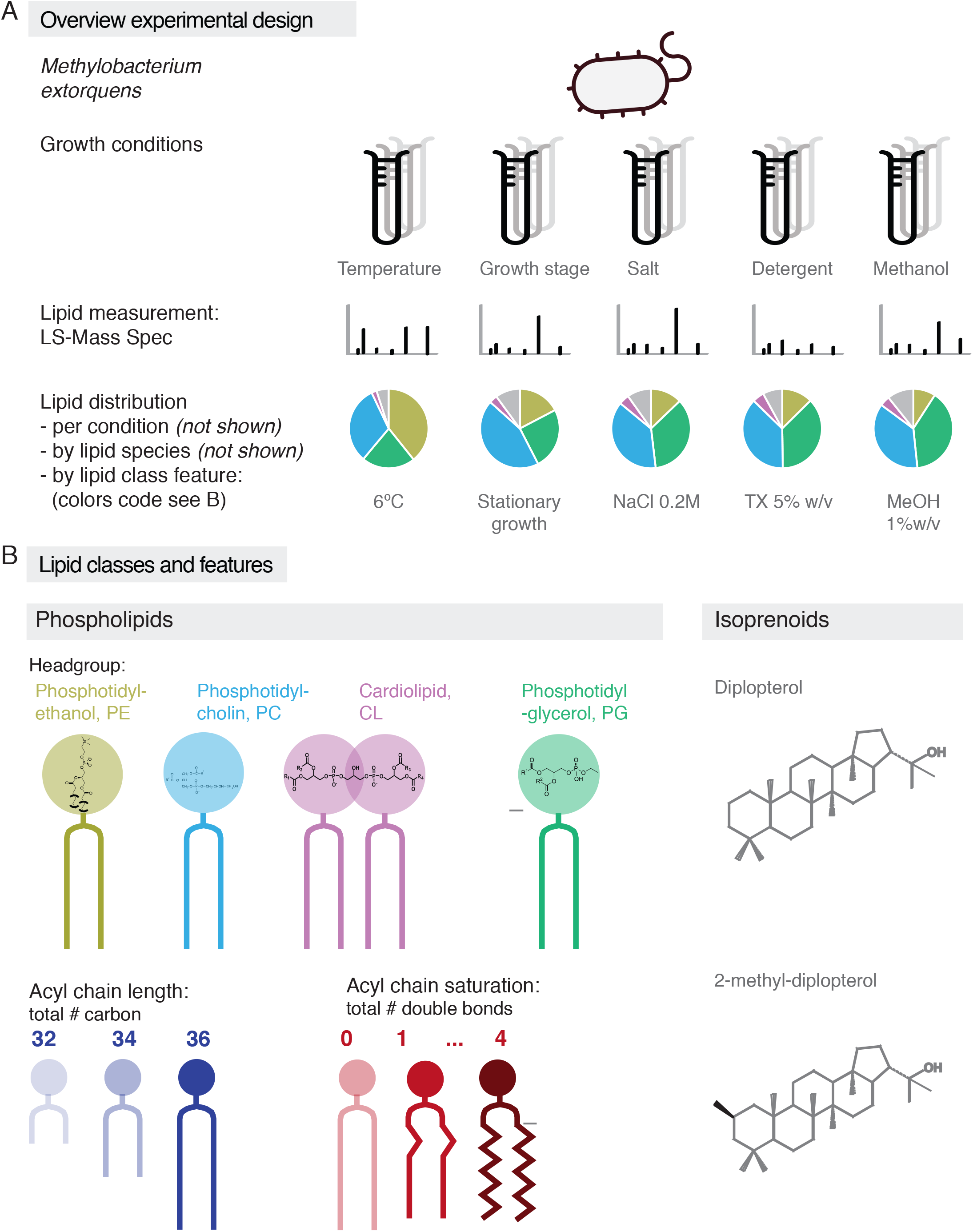
Overview of experimental outline and lipids analysed in this study. **(A)** Overview of experimental design and exemplary data obtained. **(B)** Schematic of lipid structural features measured in this study: headgroups of phospholipids and isoprenoids, acyl chain length and acyl chain saturation (number of double bonds).

**Figure 2.**
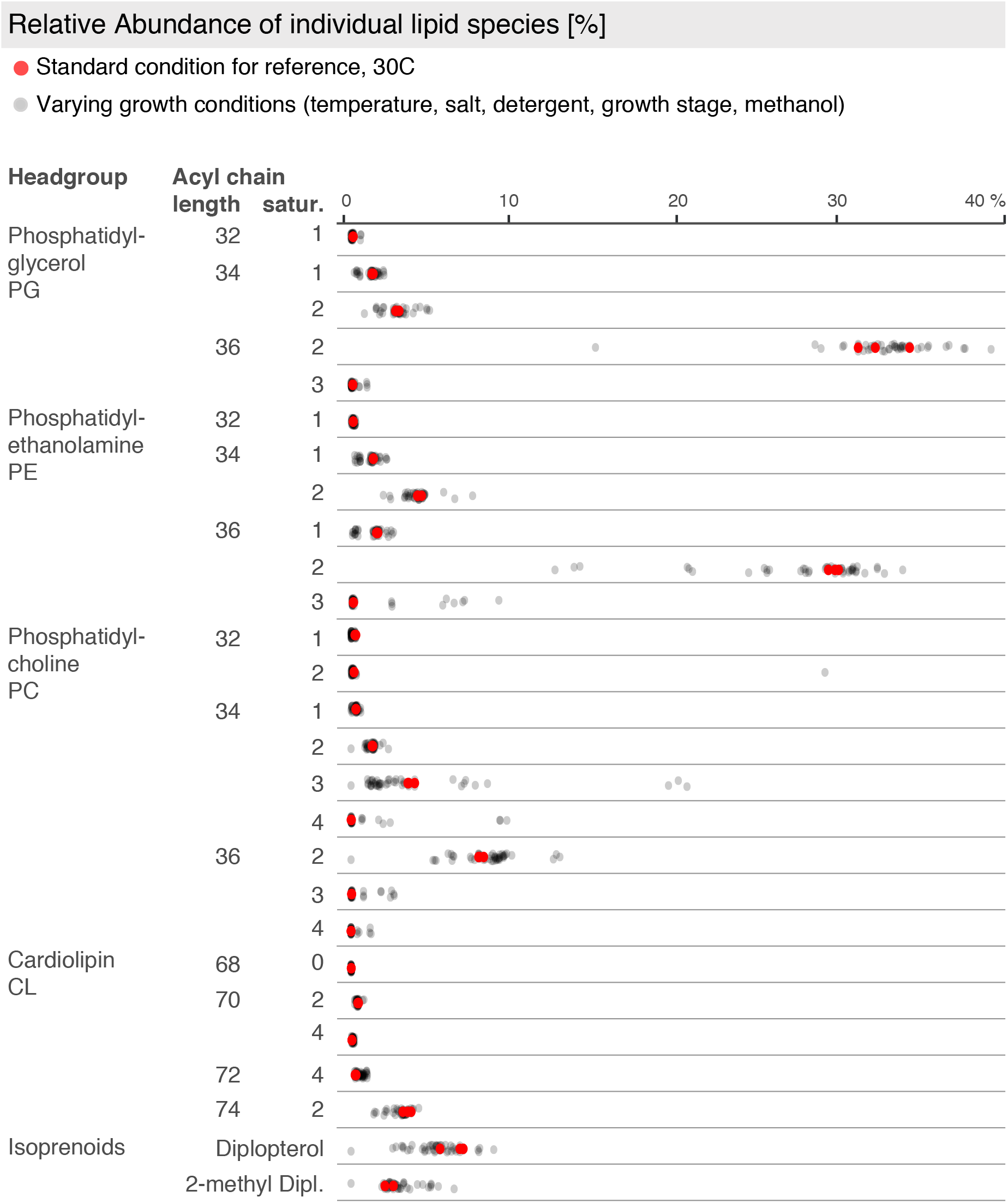
Table of lipid species abundances. Lipid species are annotated by class/headgroup (PG, PE, PC, CL or diplopterol), total acyl chain length (32 to 72), and total acyl chain saturation (number of double bonds per lipid, 0 to 4). Abundance is shown as percentage of total lipids per experiment. The range in abundance across all experimental conditions is indicated by individual grey dots, the abundances for the standard growth condition (30 °C, early exponential stage, 0.15 M sucrose) are shown in red.

It has been previously shown that *M. extorquens* produces predominately lipids with 16 and 18 carbon atoms-long acyl chains, and with up to 2 double bonds per acyl chain (Bradley et al., 2017; Wallace et al., 1990). Consistently, the major PG, PE, and PC species have cumulative chain length of 36 or 34 carbons with two double bonds per lipid (e.g. 18:1/18:1 and 18:1/16:1) and minor species with up to 4 double bonds per lipid. We also observed methylation of diplopterol (2Me-DIP) which has previously been reported in cyanobacteria and methylotrophic bacteria including *Methylobacterium* (Bradley et al., 2017; Rashby et al., 2007; Stampf et al., 1991; Summons et al., 1999). Recent work suggests that 2-methylation may be particularly prevalent among plant-associated bacteria (Ricci et al., 2014). In the following sections, we dissect the data in Figure 2 in order to reveal patterns in lipidomic variation across experimental conditions.

### 2.2 Lipidomic variation across experimental conditions

To broadly characterize the effect of environmental perturbations on membrane remodeling, we evaluated the cumulative lipid variability for each growth condition. Presently there is no gold standard for evaluating lipidomic variability. Lipid variability has previously been reported as the variance over mean of lipid abundance (Klose et al., 2012). This analysis reveals how much a lipid varies relative to its abundance, such that even a very low abundance lipid that has a small change in relation to a total lipidome could have a high variance over mean. However, in this study we aimed to characterize variations that would have an impact on the bulk adaptive properties of the membrane. Since small changes in abundance do not have a considerable effect on the bulk adaptive properties of the membrane, we focused on lipids with high absolute changes in abundance. Therefore, we calculated lipid variability as the standard deviation of abundances over a range of conditions. The total lipidomic variability was then calculated as the aggregate standard deviation of relative abundance for all individual lipid species over the range of the various conditions (e.g. temperature, growth stage, etc.) (Figure 3a). This analysis revealed that temperature has by far the largest effect on lipidomic remodeling, with more than 2-fold more variability than any other condition. Surprisingly, the smallest effects were observed with detergent challenge.

**Figure 3.**
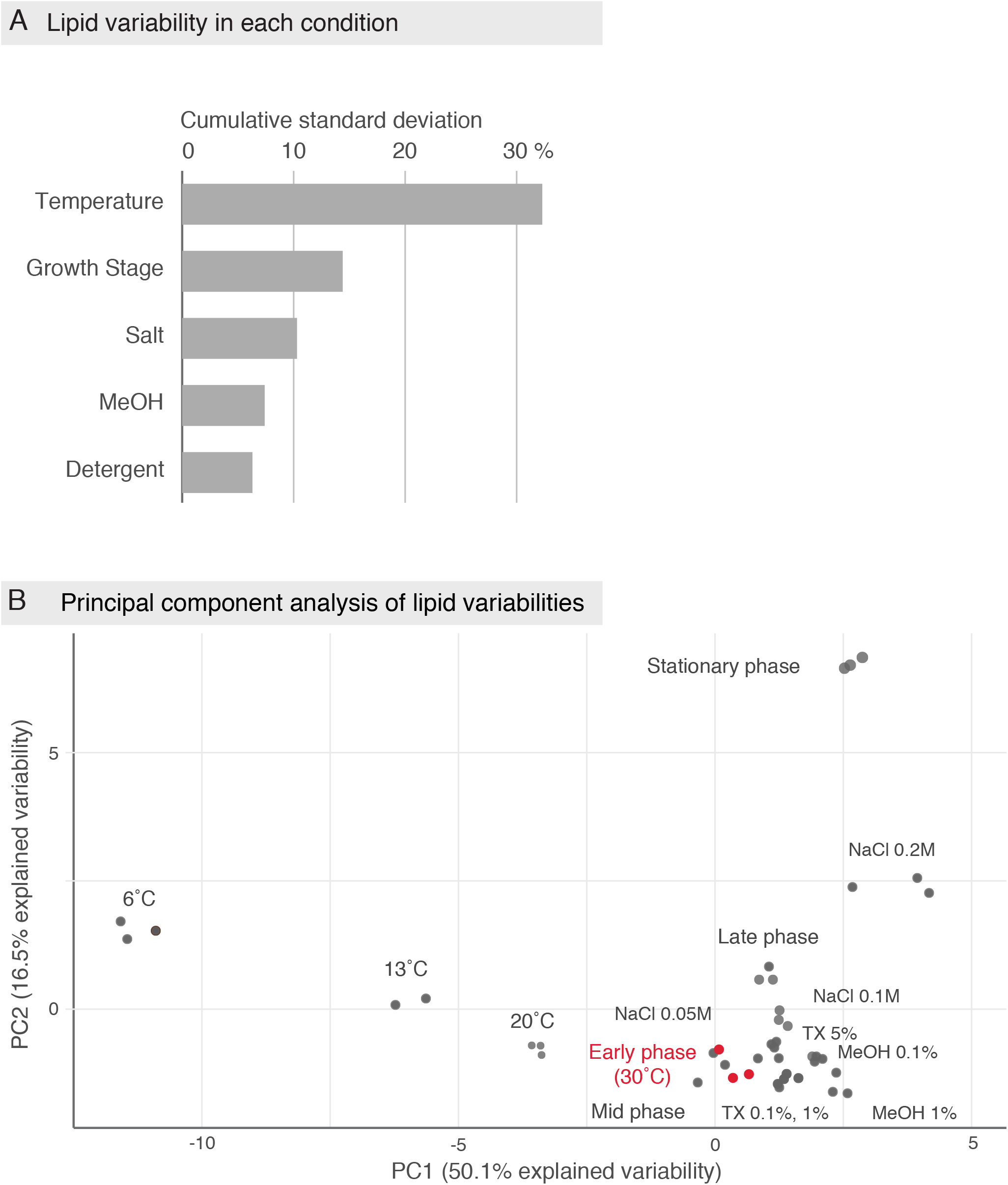
Effect of experimental conditions on lipidomic variability. **(A)** Total lipidomic variability is given as the cumulative standard deviation of individual lipid species across each set of experimental conditions. Lipid variability is highest when varying temperature, and lowest in increasing percentage of detergent. **(B)** Principle component analysis (PCA) of the lipid species abundances shows the degree of variation between subconditions. The standard growth condition (30 °C, early exponential stage, 0.15 M sucrose) is indicated in red, and all other subconditions are grey. Again, lipidomes vary greatly in changing temperatures regimes, and also when bacteria enter stationary phase growth.

To compare general lipidomic features for each individual experimental condition, we performed a principle component analysis (PCA) on lipid species abundances (Figure 3b). PCA is a method that allows for the comparison of multidimensional data by reducing variations down to two-dimensions called principle components. Distance between each point on the PCA plot is proportional to variation in lipid composition between conditions. Most conditions clustered closely with the standard growth condition, shown in red. Thus, lipid composition was not strongly affected by detergent, methanol, low salt concentrations, or during the progression through early, mid, and late growth stage. In stark contrast, bacteria grown at low temperatures, in stationary growth phase, or at high salt conditions (0.2 M NaCl) did not cluster with the standard growth condition, indicating a high extent of lipidome remodeling. Furthermore, temperature had an orthogonal effect compared to stationary growth and high salt conditions, indicating that a different set of lipidomic features is involved in responses to those conditions.

It is surprising that a membrane destabilizing detergent (Helenius, 1975) did not result in lipidomic remodeling. This could possibly be attributed to the ability of Gram-negative bacteria to resist chemicals through the robustness imparted by barrier function of lipopolysaccharide at the cell surface and active removal of toxins by multidrug transporters (Nikaido, 1996; 2003; Nikaido and Vaara, 1985; Piddock, 2006). The insensitivity of the lipidome to methanol was also unexpected. Klose et al. (2012) showed that the yeast lipidome varied greatly with differing carbon substrates based on a similar PCA analysis. Our result suggests that methanol and succinate metabolism do not require different membrane properties in *M. extorquens*. Nonetheless, the dominant effect of temperature is consistent with the large diurnal temperature variations that *M. extorquens* must adjust to in its native habitat on plant leaf surfaces and in soils. It is possible that the mechanisms of lipidome adaptation and the lipidomic composition in *M. extorquens* have evolved to be particularly well-suited for temperature changes.

### 2.3 Variation in individual lipid species

We next examined the individual lipid species involved in membrane adaptation in *M. extorquens* (Figure 4). We again calculated individual lipid species as the standard deviation of abundances for a given range of conditions (Figure 4b’). The highest abundance species generally show the largest degree of variability (Figure 4a and 4b). However, there are some low abundance species that exhibit large variations, notably lipids with 3 and 4 double bonds. An important point of emphasis is that low abundance or undetectable species at standard growth conditions can become major components under certain conditions, playing a crucial role in adaptation. The most striking result is that the majority of lipid species do not vary substantially under any conditions, suggesting that only a small fraction of the lipidome is required for adaptive membranes.

**Figure 4.**
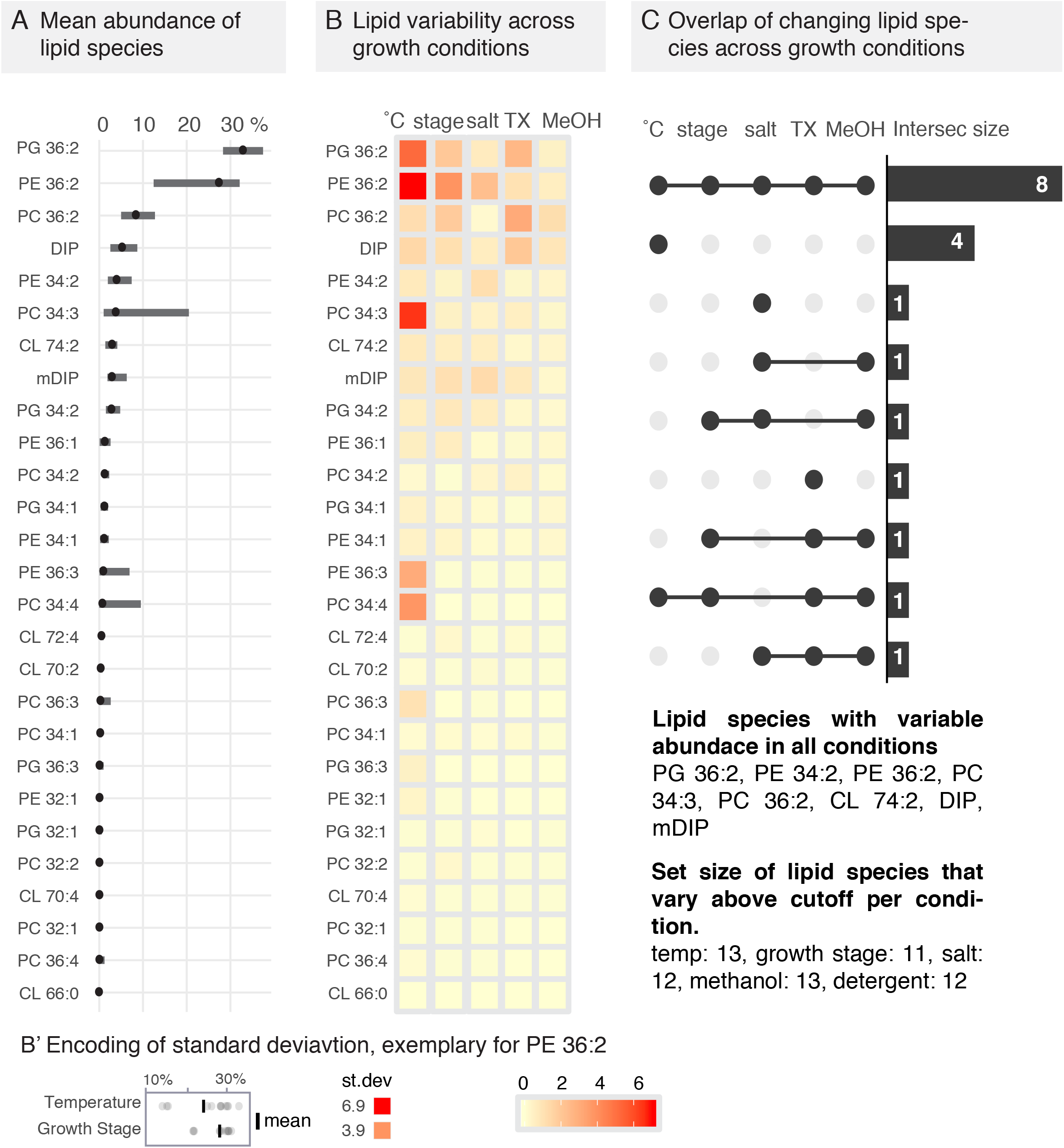
Variability of individual lipid species across experimental conditions. **(A)** Average abundances of lipid species for all conditions sorted from highest (PG 36:2) to lowest (CL 64:0) lipid species. Mean and range are shown for each lipid. **(B)** Heat map of lipid species variability across experimental conditions, high variability is indicated in red, no variability in light yellow. **(B’)** Exemplary calculation of standard deviation. **(C)** UpSetR plot (Conway et al., 2017) showing the overlap of lipids adapting their amounts across the experimental conditions. Eight lipids vary their abundance in all experimental conditions, four lipids vary abundance specifically in changing temperatures. Stage: bacterial growth stage/phase, TX: varying detergent concentrations.

We then asked how many and which lipid species are highly variable for each set of conditions. We defined the highly variable lipid species as those that account for 90% of the total lipidomic variability within any given condition. By this measure we observed that 10-13 lipids are highly variable over any given condition (Figure 4c), meaning that only around 30% of the lipidome is involved in remodeling during adaptation. Of these 10-13 highly variable lipids, 8 are highly variable over all of the conditions tested. These results revealed that only a fraction of the lipidome is involved in adaptive remodeling and point towards the minimum complexity of lipid species required for membrane adaptation. In principle, it is possible that as few as 8 lipid species could support adaptation over the range of experimental conditions examined in this study.

### 2.4 Variation of lipid structural features

Membrane biophysical properties are modulated by changes in the abundance of lipid structural features. For instance, increasing the number of double bonds per lipid reduces membrane viscosity during adaptation to decreasing temperature, whereas various lipid headgroups can impart curvature or surface charge. We analyzed how the various structural features of *M. extorquens* lipids are modulated to glean insight into their sense and response mechanisms of membrane adaptation.

Headgroup features were differently regulated, as PC, PE and PG were relatively plastic, whereas CL and diplopterols are relatively invariant (Figure 5a). For acyl chains, most of the variability is observed for lipids with 2 and 3 double bonds and with total chain length of 34-36 carbons (Figure 5b). When we group acyl chain features separately for each headgroup (e.g. PC-specific unsaturation) PC shows the largest variability for both unsaturation and chain length, whereas PG acyl chain features are nearly invariant. This shows that acyl chain features are regulated independently within each headgroup and provides a starting recipe for designing a responsive synthetic lipidome.

**Figure 5.**
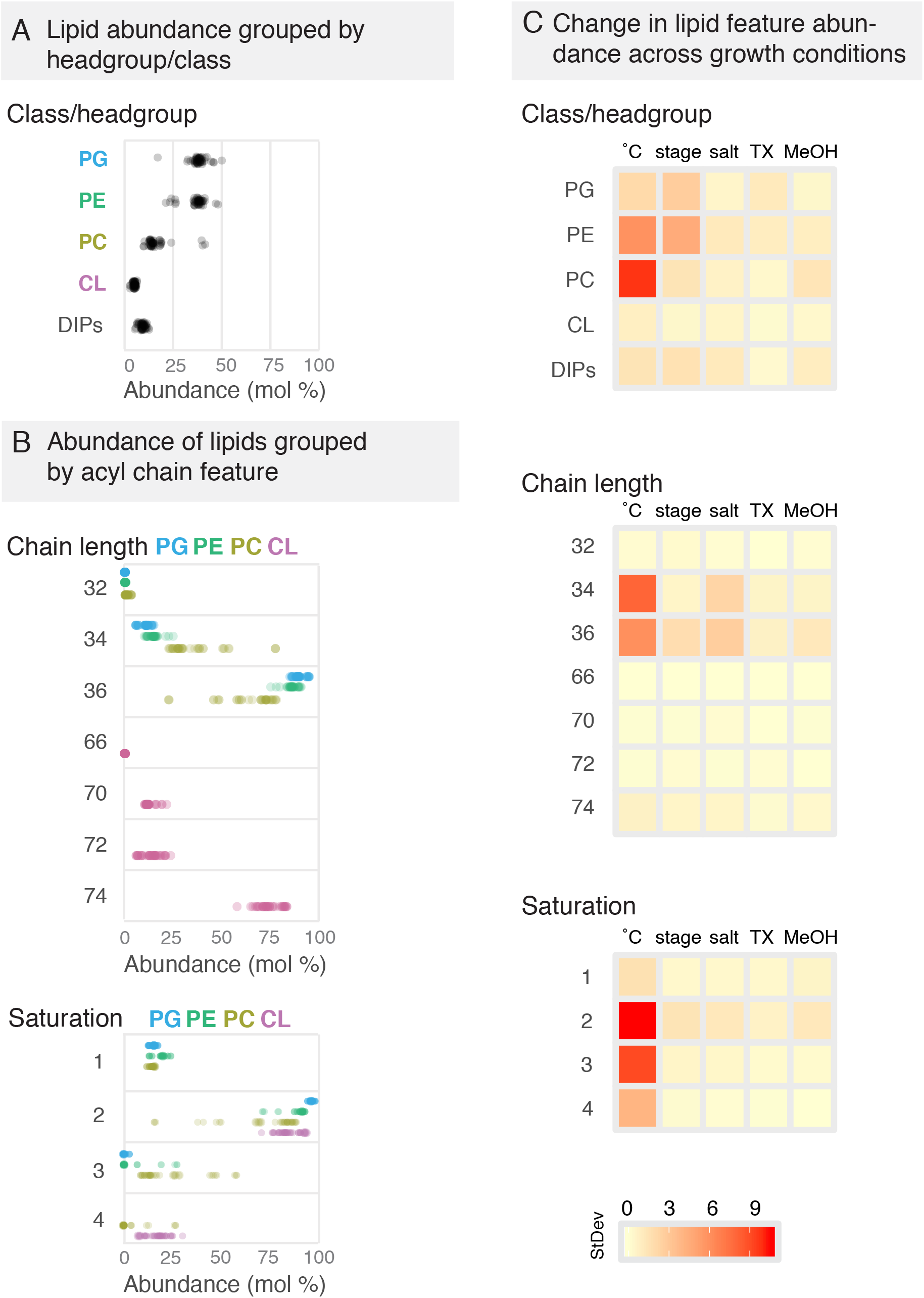
Variability of phospholipid structural features across experimental conditions. **(A)** Abundances of lipids grouped by lipid class shows overall low variability, and lowest variability of CL. **(B)** Abundances of phospholipid headgroup binned by acyl chain length and acyl chain saturation respectively reveals group specific differences in acyl chain remodeling. (headgroups: blue = PG, green = PE, olive = PC, magenta = CL). **(A), (B)** each data point shows lipid abundance from individual biological replicates. **(C)** Variability of lipid features plotted for each set of experimental conditions. Stage: bacterial growth stage/phase, TX: varying detergent concentrations.

To assess how much different features varied for each experimental condition we calculated the variance of lipid features for each range of conditions as depicted by the heat map in Figure 5c. The highest variation in lipid classes is observed over temperature and growth stage. Interestingly, total lipid unsaturation varies over temperature but is relatively invariant over all other conditions. The most variation in total lipid chain length varies with temperature and salt concentration. These results reveal that membrane adaptation of total lipid features exhibit different patterns in response to different experimental conditions. This shows that the lipidomic adaptation mechanism is capable of unique, orthogonal responses to different perturbations.

### 2.5 Progressive remodeling of lipids across experimental conditions

Assessing lipidomic data for each step within each condition allows us to compare lipids as we go stepwise through the changes. For instance, how does the lipidome compare when the growth temperature is lowered from 30 to 20 to 13 and to 4 °C, or as cells progress from early to mid, to late and finally to stationary growth phase? We again consider lipids grouped by their features as this allowed us to monitor structural variations that directly impact the membrane’s biophysical properties (Figure 6; Figure 6-Figure supplement 1-4).

**Figure 6.**
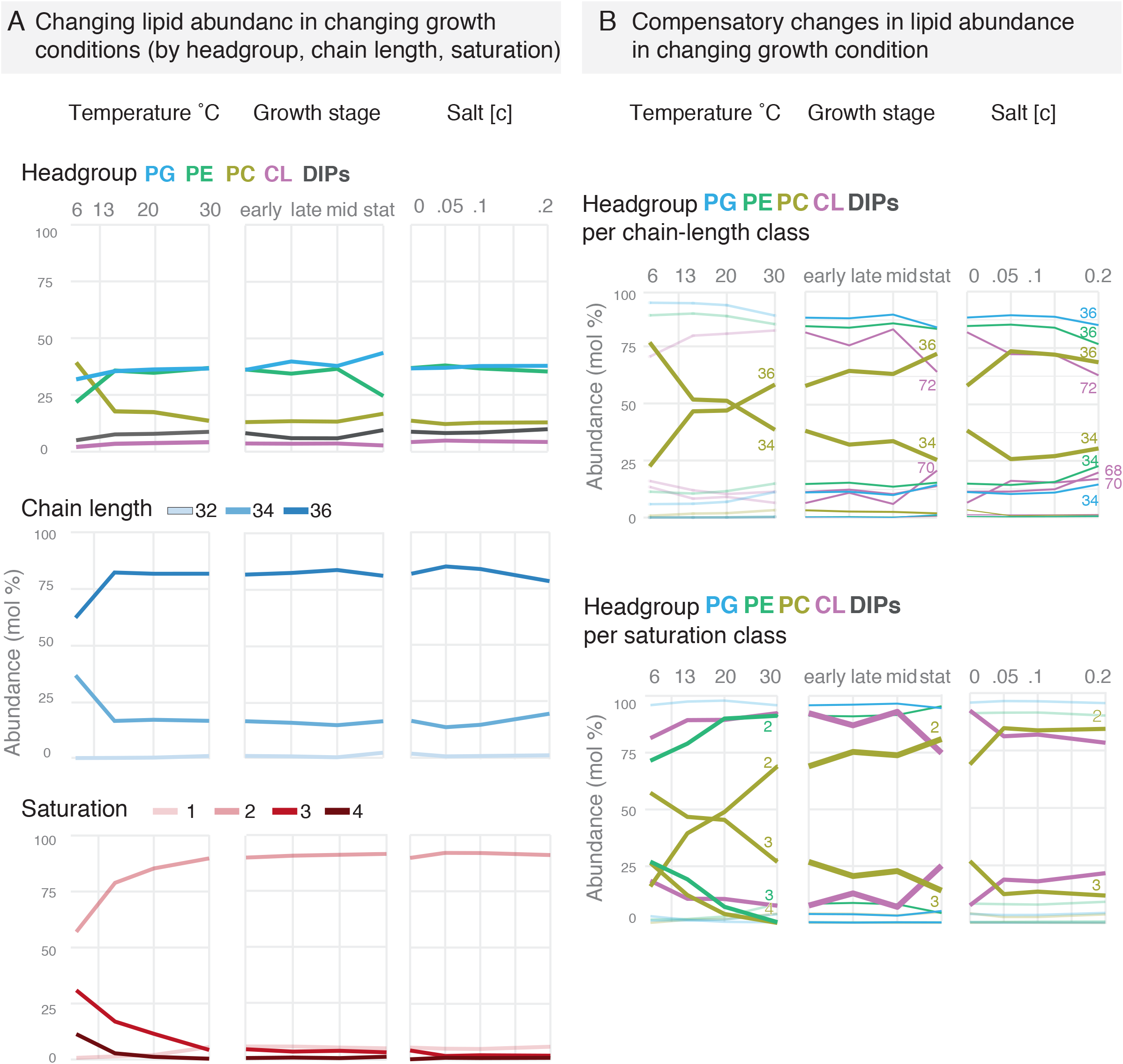
Step-wise adaptation of lipids by feature at changing environmental growth conditions. **(A)** Abundances of lipids grouped by class, acyl chain length and acyl chain saturation when changing growth temperature, probing at different growth stages, and varying salt (NaCl) concentration of the growth media. **(B)** Abundances of phospholipids grouped by acyl chain features, with headgroups indicated by color (blue: PG, green: PE, olive: PC, magenta: CL). This reveals compensatory effects in lipid adaptability: while abundance of PC with 2 double bonds increases with increasing temperatures, PC with 3 double bonds decreases from 6 to 30 °C. In each panel, the most variable features are highlighted in bold.

#### Class

Lipid class abundances are relatively invariant over most conditions (Figure 6). PC abundance increases with decreasing temperature, with a pronounced change at 6 °C. PE abundance increases considerably when cells are in stationary growth stage. These results suggest that lipid class distribution is only adjusted under extreme conditions.

#### Double Bonds

The total number of double bonds per lipid varies substantially with temperature and is nearly invariant for other conditions (Figure 6a). As temperature decreases the abundances of DB3 and DB4 lipids increase, while abundances of DB1 and DB2 decrease. This contrasts with other organisms such as yeast where primarily DB1 and DB2 are adjusted with temperature change (Klose et al., 2012). It is possible that having additional double bonds enhances Methylobacterium’s ability to adapt to broad temperatures that they experience on plant surfaces and in soils.

The variation of double bonds per lipid within each headgroup class is not evenly distributed across classes (Figure 6b). For instance, for varying temperatures PC shows the highest remodeling whereas PG is nearly invariant. Additionally, for varying salt and growth stages where total unsaturation has little or negligible variability, class-specific PC-DB and CL-DB vary considerably. These variations in PC-DB and CL-DB are nearly equal and opposite resulting in no apparent change in total DB content.

#### Chain length

Total chain length is highly conserved across all conditions, and only varies substantially at 6 °C (Figure 6a). Since total chain length has a large effect on membrane thickness, which in turn must be matched with membrane proteins transmembrane domain length, it is not surprising that this feature is so invariant. The large change in total CL at extreme low temperature suggests that the membrane must take extreme measures to adapt as it approaches the end of the range of viable growth temperature.

Similar to unsaturation, chain length within each headgroup class shows much higher variability than total chain length (Figure 6b). For instance, from 30 to 20 °C total chain length remains constant while PC chain length decreases and is compensated for by increased chain length in PG, PE and CL. Similar remodeling is also observed for growth stage and salt concentration. Additionally, PC exhibits the highest chain length variation.

#### Diplopterol 2-methylation

A particularly striking variation was observed for the sterol analogue diplopterol and its 2-methylated derivative (Figure 7). Diplopterol is the major hopanoid in *M. extorquens* and is localized in the outer membrane (Hancock and Williams, 1986; Sáenz et al., 2015). We previously demonstrated that diplopterol has a sterol-like ability to modulate the order of saturated lipids such as lipopolysaccharides in the bacterial outer membrane (Sáenz et al., 2015; 2012). Subsequent work demonstrated that 2-methylation of hopanoids increases their membrane rigidifying effect (Wu et al., 2015). Therefore, variations in diplopterol 2-methylation could play a role in the adaptation of the outer membrane. While total diplopterol abundances (diplopterol + 2-methyl-diplopterol) did not vary substantially under any conditions, 2-methylation of diplopterol is highly variable in all conditions except methanol and detergent (Figure 7a and b). There is a progressive increase in 2-methyl-diplopterol abundance with decreasing temperature and increasing salt concentration (Figure 7c), consistent with transcriptional regulation of this process by global stress response pathways (Kulkarni et al., 2013). These results show for the first time that 2-methylation is dynamically regulated in response to varying growth conditions and is important for outer membrane adaptation.

**Figure 7.**
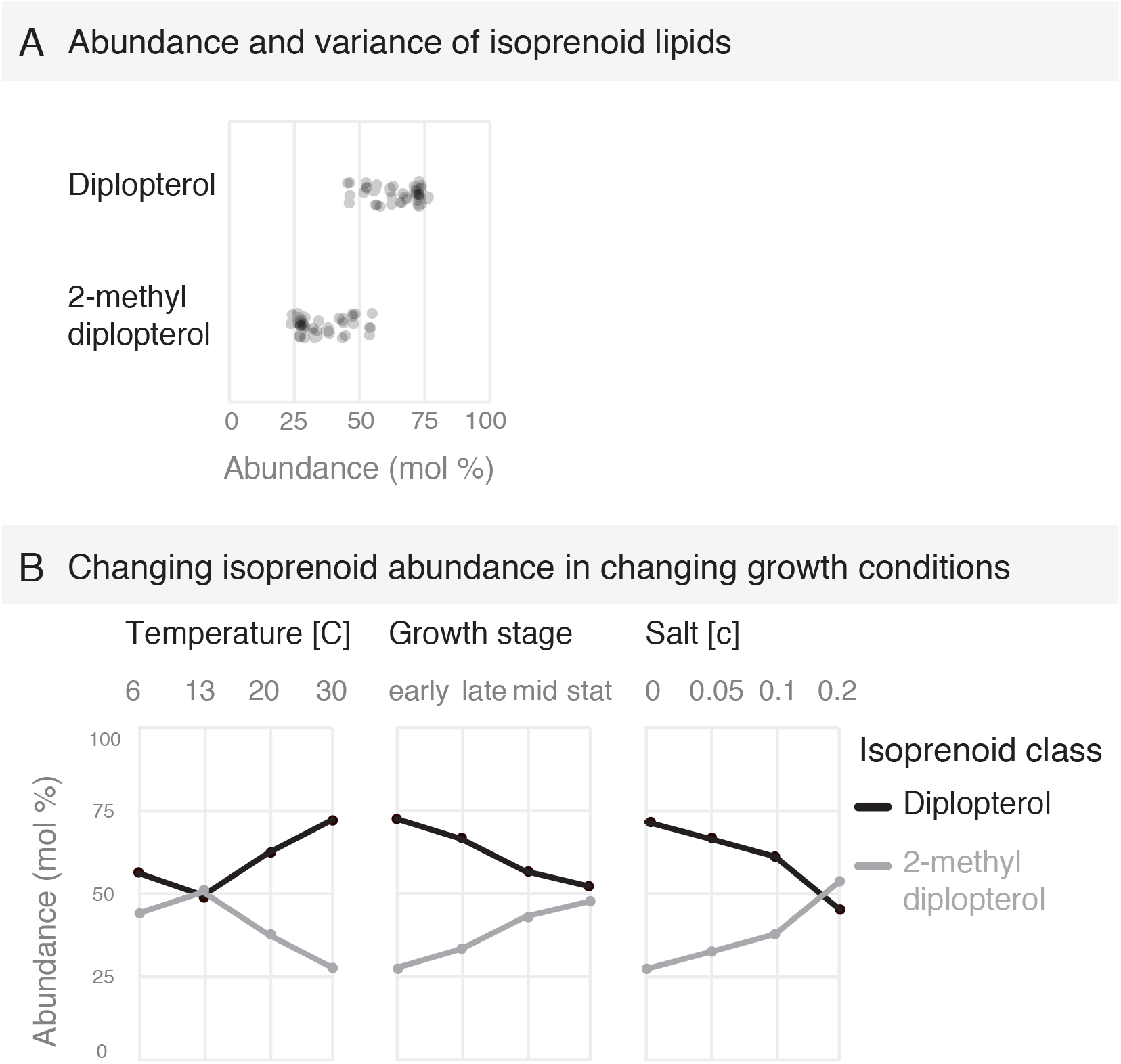
Diplopterol (hopanoid) methylation. **(A)** Abundances of diplopterol species across growth conditions. **(B)** Step-wise adaptation of diplopterols when changing temperature, growth phase, and salt condition.

Our observations highlight three characteristics of structural lipid adaptation. First, we observed that adaptivity of features over a particular range of conditions (e.g. temperature) does not involve a monotonic pattern of feature remodeling, but rather can involve different features over intervals of a given condition. Second, certain features such as acyl chain unsaturation and diplopterol 2-methylation are highly variable and are remodeled over a broad range of conditions. However, other features such as lipid class and acyl chain length are relatively invariant except under extreme conditions of low temperature and stationary growth stage, suggesting that chain length and headgroup distribution have multiple constraints (e.g. maintaining membrane thickness or surface charge). Notably, even the tightly constrained features are adjusted under extreme perturbations. Third, acyl chain features are regulated independently within each headgroup class. Thus, even when total acyl chain unsaturation and length appears invariant, there can be extensive acyl chain remodeling within specific headgroup classes. Such class-specific remodeling of unsaturation may result in adaptation that is localized to the inner or outer membrane, or to specific regions within each membrane, such as highly curved regions at the poles, one particular membrane leaflet, lipids that are closely associated with protein transmembrane domains, or potentially lateral membrane domains. On the other hand, constraining total unsaturation while internally shifting double bonds from PC to CL might be a means of varying intrinsic curvature while maintaining total membrane viscosity.

### Summary and Outlook

In this study, we systematically characterized the lipidome of *M. extorquens* over a range of growth conditions to gain insights into membrane homeostatic adaptation. Countless studies have addressed phospholipid acyl chain composition (e.g. as measured by GC or GC-MS) and have revealed an important role of membrane lipid composition to various phenotypic aspects. However, phospholipid headgroups and acyl chains jointly determine membrane properties. Using shotgun lipidomics, we provide a near-complete lipidome down to the species level. This allowed us to map how structural features of lipids (e.g. head group, saturation, chain length) are recombined to achieve adaptation. We demonstrated that membranes composed of only a few lipid species are capable of adapting specifically to a broad range of conditions, in particular temperature. Additionally, by assessing how lipid structural features are recombined through adaptation, we reveal that even a simple lipidome exhibits complex patterns of remodelling that were previously undefined. We propose that it is essential to assess lipidomic data in terms of the composite structural features conferred by binned combinations of lipids, since the combination of these structural features ultimately dictates the physical state and functionality of membranes.

We probed the lipidome composition across a broad range of viable conditions spanning temperature, hyperosmotic and detergent stress, carbon sources, and cell density. PCA analysis revealed that most lipidomes are highly similar across conditions (Figure 3). This demonstrates that the lipidome is robust and that there are strong constraints on lipidome composition and remodelling. While individual lipid species do adapt to changing conditions, eight lipid species are variable in all experimental regimes (Figure 4) and the majority of lipids remained surprisingly stable. One of the outstanding conundrums in membrane biology is to understand why there are so many lipids (Dowhan, 1997; 2017), even a relatively simple organism like *E. coli* has over 100 phospholipid species (Jeucken et al., 2019). Recapitulating such a complex system from first principles seems daunting. However, the lipidome of *M. extorquens* demonstrates that a synthetic membrane capable of supporting life over a broad range of environmental conditions could be constructed with a relatively small number of lipid species.

Lipidome adaptation is also constrained by structural elements of individual lipids. Instead of analysing lipidomes purely by lipid species, it is therefore critical to assess global structural composition. For example, total phospholipid chain length, that influences membrane thickness, is a highly conserved feature that only varies under extreme conditions such as low temperature (Jensen and Mouritsen, 2004). Interestingly, phospholipid acyl chain length was highly conserved globally, but variable within certain headgroup classes. Other constraints do not have such clear implications. For instance, phospholipid headgroup composition only varies at low temperature or in stationary growth phase, possibly suggesting conservation of membrane surface charge as a key constraint. Also, the acyl chain composition among phospholipids with PG headgroup is nearly invariant across all conditions, suggesting that there could be biochemical constraints on PG remodelling. Low PG remodelling has also been observed in cyanobacteria (Pittera et al., 2018), showing that this constraint is not limited to *M. extorquens*. While the physicochemical and physiological basis for these observations remains to be fully explored, the patterns and constraints in lipidome remodelling that we report here serve as a resource for designing a minimal adaptive membrane and highlight the need for multi-level analysis of lipidome data.

Among all phospholipid classes, we observed that phosphotidylcholine (PC) showed the highest degree of acyl chain remodelling. This is particularly interesting with regard to bacterial-host interactions. While relatively few bacteria synthesize PC, the majority of studied bacteria associated with a eukaryotic host, including many pathogens, contain PC in their membranes (Aktas et al., 2010; Geiger et al., 2013). Synthesis of PC was previously shown to be essential for host colonization (Minder et al., 2001) and for pathogen virulence (Conde-Alvarez et al., 2006; Wessel et al., 2006). The specific physiological role of bacterial PC, however, is still unclear. Our results that PC accounts for most acyl chain remodelling in *M. extorquens* could suggest a pivotal role of membrane adaptation through PC remodelling for bacterial-host interactions.

Understanding how *M. extorquens* adapts with so few lipids, and even fewer that are responsive, remains an open challenge. In particular, given that lipid remodelling did not involve a monotonic change, but rather seemed to involve different features varying over different intervals of a given range of conditions, such as temperature. Critical to understanding the mechanisms underlying lipidome adaptation are the sense and response pathways that regulate the remodelling of lipid structural features. It has emerged in recent years that there can be dedicated membrane sensors, which are involved in regulating lipid biosynthesis pathways in response to perturbed membrane properties (Ernst et al., 2016; Puth et al., 2015; Saita and de Mendoza, 2015). Recently, it was demonstrated that in addition to membrane thickness and viscosity, some membrane sensors are poised to detect changes in lateral tension in the bilayer (Ballweg et al., 2019), suggesting a diverse range of possible inputs to lipidome adaptation. Building on the lipidomic resource presented here, *M. extorquens* is a promising model organism to explore minimal mechanisms for cell membrane adaptation. The adaptive lipidome of *M. extorquens* provides a critical resource as a starting point to elucidate the regulatory systems underlying membrane adaptation.

## Acknowledgements

The authors wish to thank Kai Simons and Robert Ernst for advice, members of Sáenz group for discussions, Michael Schlierf, Ilya Levental, Alex Bradley, Carl Modes, and Christoph Zechner for manuscript comments, and Lipotype GmbH.

## Author contributions

Sandra Rizk and Daniel Grosser performed bacterial growth experiments. Oksana Lavrynenko performed MS analysis. Grzegorz Chwastek contributed to writing, editing and data analysis. Michał Surma performed MS data analysis and editing. Magda Rucińska performed bioinformatics analysis. Helena Jambor conceived and designed the figures. James Sáenz conceived the project, obtained funding, supervised the project, and wrote the manuscript. All authors read, edited and approved the final manuscript.

## Methods

### Media, growth conditions

For all cultivations a defined media, here referred to as Hypho-TMM, was used (Delaney et al., 2013). All liquid cultures were grown at 30 °C and pH 7 with shaking at 150 rpm unless stated otherwise. For all experiments we used a strain of *M. extorquens PA1* with the cellulose synthase genes knocked out to minimize clumping and improve cell density measurements (Delaney et al., 2013). Cultures were started from a glycerol stock stored at −80 °C and were plated on Hypho-TMM agarose plate and grown at 30 °C. A liquid pre-culture was set up by sterile picking a single colony and transferring it to 5 ml of Hypho-TMM. This pre-culture was grown over night to near stationary phase. The experimental culture was started by inoculation to OD_600 nm_ 0.02 with pre-culture and grown until a final OD_600 nm_ of 0.2 – 0.3 (early log-phase) was reached. Several conditions were tested including the influence of salinity, carbon source (MeOH and succinate), detergent (up to 5% Triton X-100) and temperature (6 – 30 °C). Cells were also harvested at mid-, late- and stationary growth stages. A defined volume representing 6 OD units (1 OD unit = 1 mL of OD 1) was sampled and pelleted. The harvested cells were washed twice with fresh media. The pellet was snap frozen in liquid nitrogen and stored until further processing at −20 °C.

#### Hypho-TMM recipe

1.45 mM K_2_HPO_4_; 1.88 mM NaH_2_PO_4_; 45.6 µM sodium citrate; 1.2 µM ZnSO_4_; 1 µM MnCl_2_; 18 µM FeSO_4_; 2 µM (NH_4_)_6_Mo_7_O_24_; 1 µM CuSO_4_; 2 µM CoCl_2_; Na_2_WO_4_; 0.5 mM MgCl_2_;

8 mM (NH_4_)_2_SO_4_; 20 µM CaCl_2_

30 mM PIPES pH 7

15 mM succinate unless stated otherwise

### Shotgun mass spectrometry

For the lipidomic analysis by mass spectrometry the lipids of harvested cells were extracted twice by the method of Blight and Dyer (Bligh and Dyer, 1959). The total lipid extracts representing 6 OD units of *M. extorquens* were analysed by quantitative shotgun lipidomics (Ejsing et al., 2009) adapted for the quantification of hopanoids (diplopterol and 2-methyl-diplopterol).

#### Measurement of phospholipids

For absolute quantification extracts were diluted with MS mix (7.5 mM ammonium acetate in isopropanol/methanol/chloroform 4:2:1) containing the 0.5 µM PC (34:0), PE (34:0), PG (34:0) mixture and 0.05 µM CL (57:4). MS spectra were acquired on Q Exactive tandem mass spectrometer (Thermo Fisher Scientific, Bremen, Germany) equipped with a robotic nanoflow ion source TriVersa NanoMate (Advion BioSciences, Ithaca NY). Spectra were acquired in negative MS mode with the mass resolution of R_m/z 400_ = 280000 with AGC target value of 10^6^ and maximum injection time of 1 s in the range of masses 400 – 1000 m/z. Raw data were processed using LipidXplorer software (Herzog et al., 2012; 2011). Based on high accuracy and resolution masses, species belonging to 4 phospholipid classes including phosphatidylcholine (PC), detected as an acetate adduct, phosphatidylethanolamine (PE), phosphatidylglycerol (PG), both detected as deprotonated anions, and cardiolipin (CL), detected as a doubly deprotonated anion, were identified.

#### Measurement of diplopterol

The extract was dried down and re-dissolved in MS mix (7.5 mM ammonium acetate in isopropanol/methanol/chloroform 4:2:1) containing the 0.13 mM cholesterol D7 standard. MS spectra were acquired on Q Exactive tandem mass spectrometer (Thermo Fisher Scientific, Bremen, Germany) equipped with a robotic nanoflow ion source TriVersa NanoMate (Advion BioSciences, Ithaca NY). Spectra were acquired in positive MS mode with the mass resolution of R_m/z 400_ = 140000 with AGC target value of 10^6^ and maximum injection time of 1 s in the range of masses 360 – 455 m/z. Raw data were processed using LipidXplorer software. Diplopterol and 2-methyl-diplopterol were detected as M -[H_2_O] + H^+^ … and cholesterol D7 as M - [H_2_O] + H^+^. Absolute concentrations of diplopterol and methyl-diplopterol were determined using calibration curves and response factors established alongside with cholesterol D7 and purified diplopterol mixed at various ratios.

### Data analysis

Lipid abundances (mol% of total) of biological triplicates for all conditions are provided in the supplementary materials. One of the replicates at 13 °C and at TX-100 1% yielded erroneous abundance data due to a problem with the internal standard in those replicates and we have therefore excluded them from our analyses.

Variation of lipid species across experimental conditions was calculated by taking the standard deviation of lipid abundances for replicates over a given range of conditions (e.g. temperature, salt, etc.). Total lipidomic variation was calculated as the cumulative variation of individual lipid species for a given range of conditions.

Visualization of lipidome variation for all conditions was investigated with principle component analysis (PCA). PCA was performed by using the R prcomp function in RStudio with default parameters, where the calculation is carried out by a singular value decomposition of the (centered and scaled) data matrix. Visualization in single plots performed using the ggbiplot function. PCA allows the identification of latent variables (principal components) in the data based on 14 observed variables. The plot is based on principal components 1 and 2, which explain 50.1% and 16.5% of the total variance of the data, respectively.

Variations in abundance of lipid features were analysed by binning lipid species according to structural features including total chain length, saturation, class (e.g. phospholipid headgroup and diplopterols), and 2-methylation of diplopterol. Acyl chain length and saturation were binned for all phospholipids (global acyl chain features), and separately within each headgroup (headgroup specific acyl chain features).

---

**Figure 1 – supplement 1.**
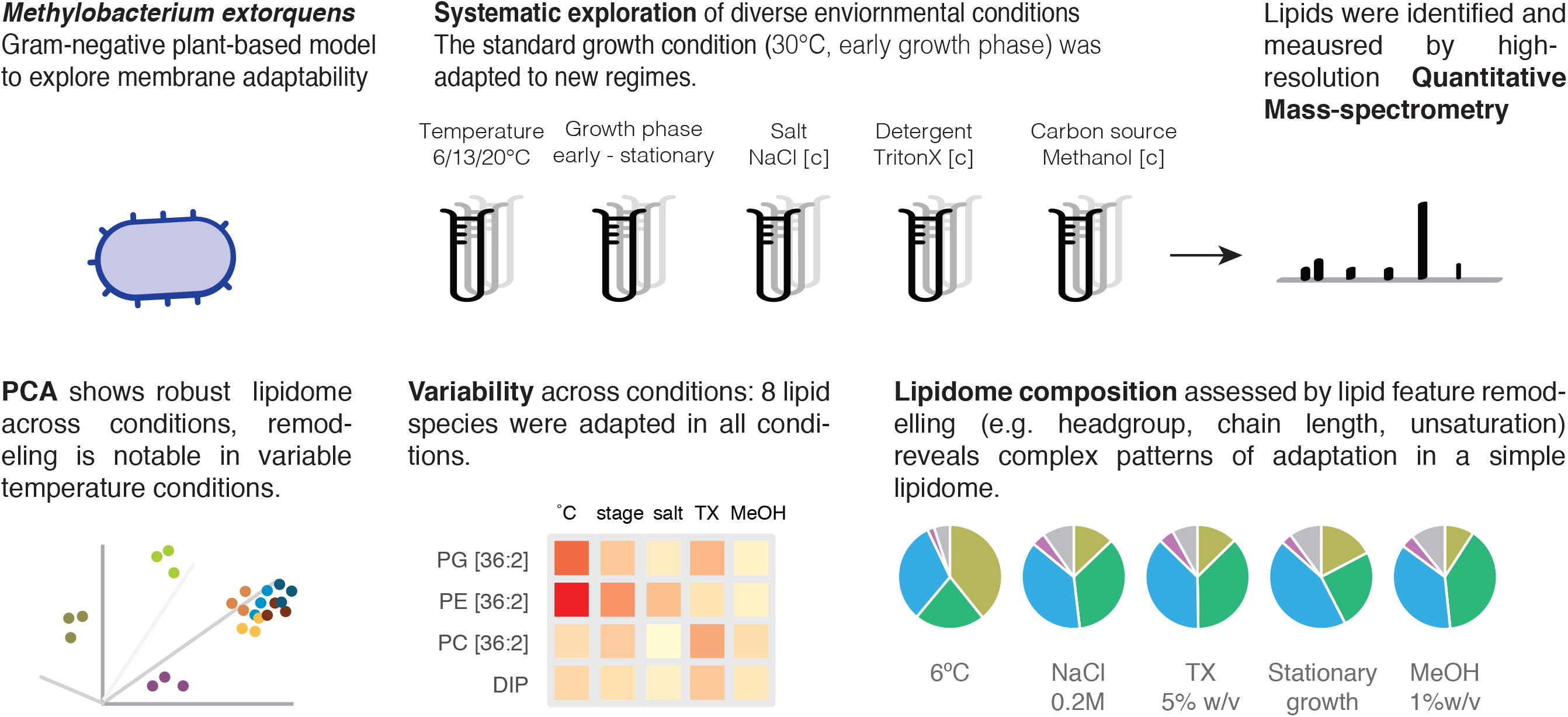
Bacterial growth rates (Herbert et al., 1956) for all experimental conditions.

**Figure 1 – Supplement 2.**
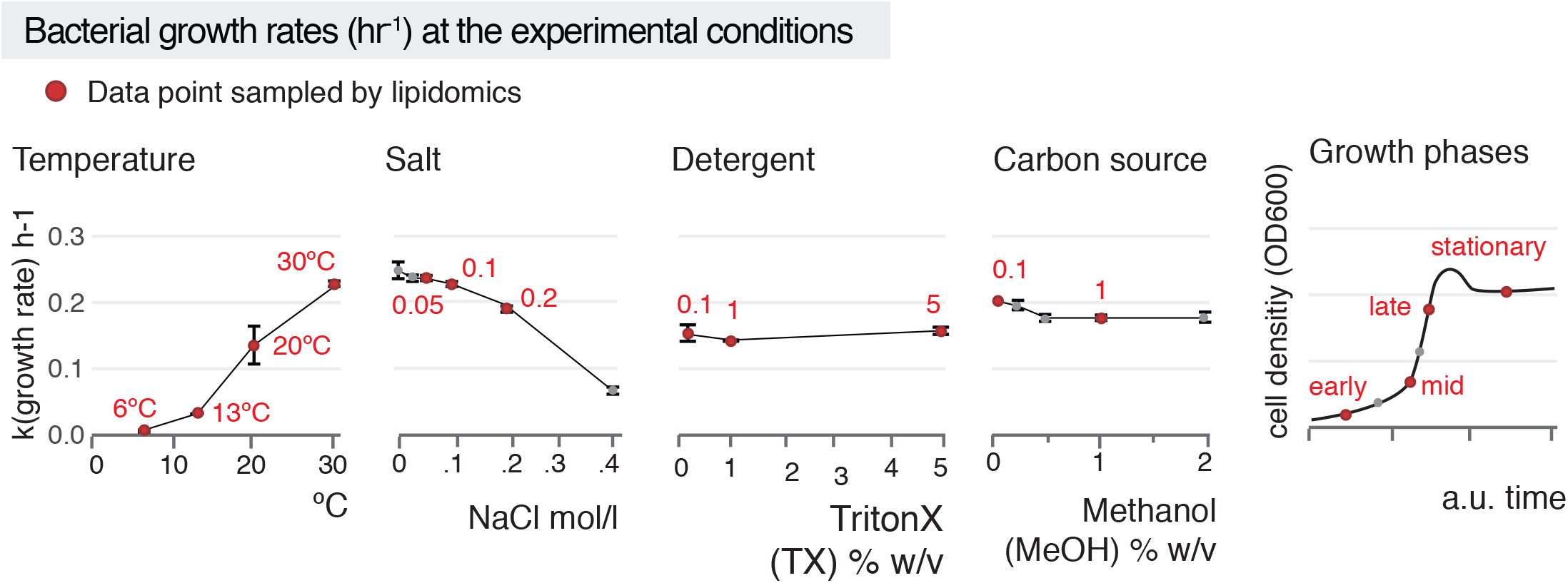
Bacterial growth curves and sampling points for lipidomic analysis. Chart for growth phases is a conceptual drawing.

**Figure 6 – Supplement 1.**
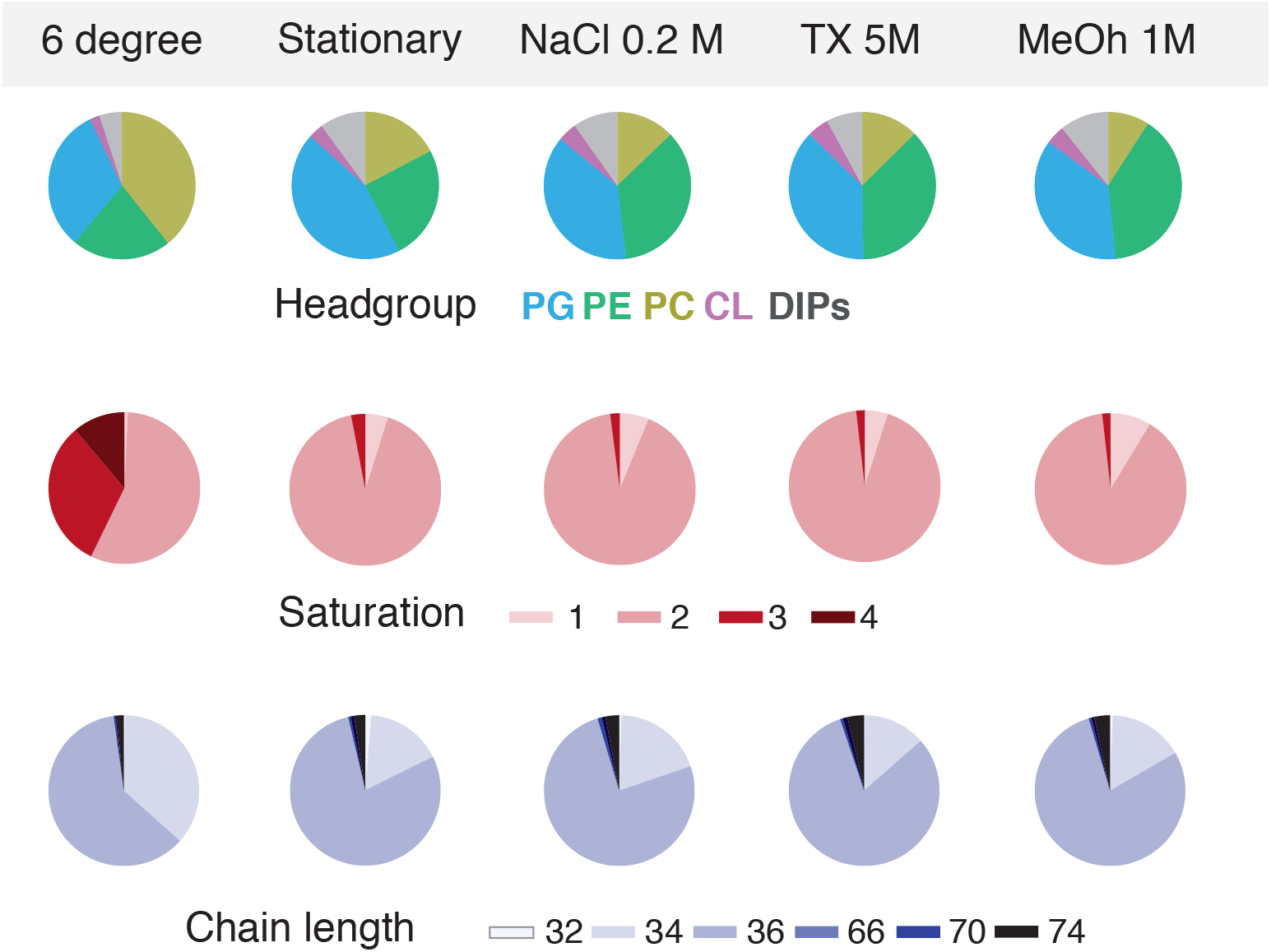
Overview of compositional changes across exemplary experimental conditions reveals the lipidome is robust in most growth conditions except in drastic temperature changes when the lipidome is markedly different.

**Figure 6 – Supplement 2.**
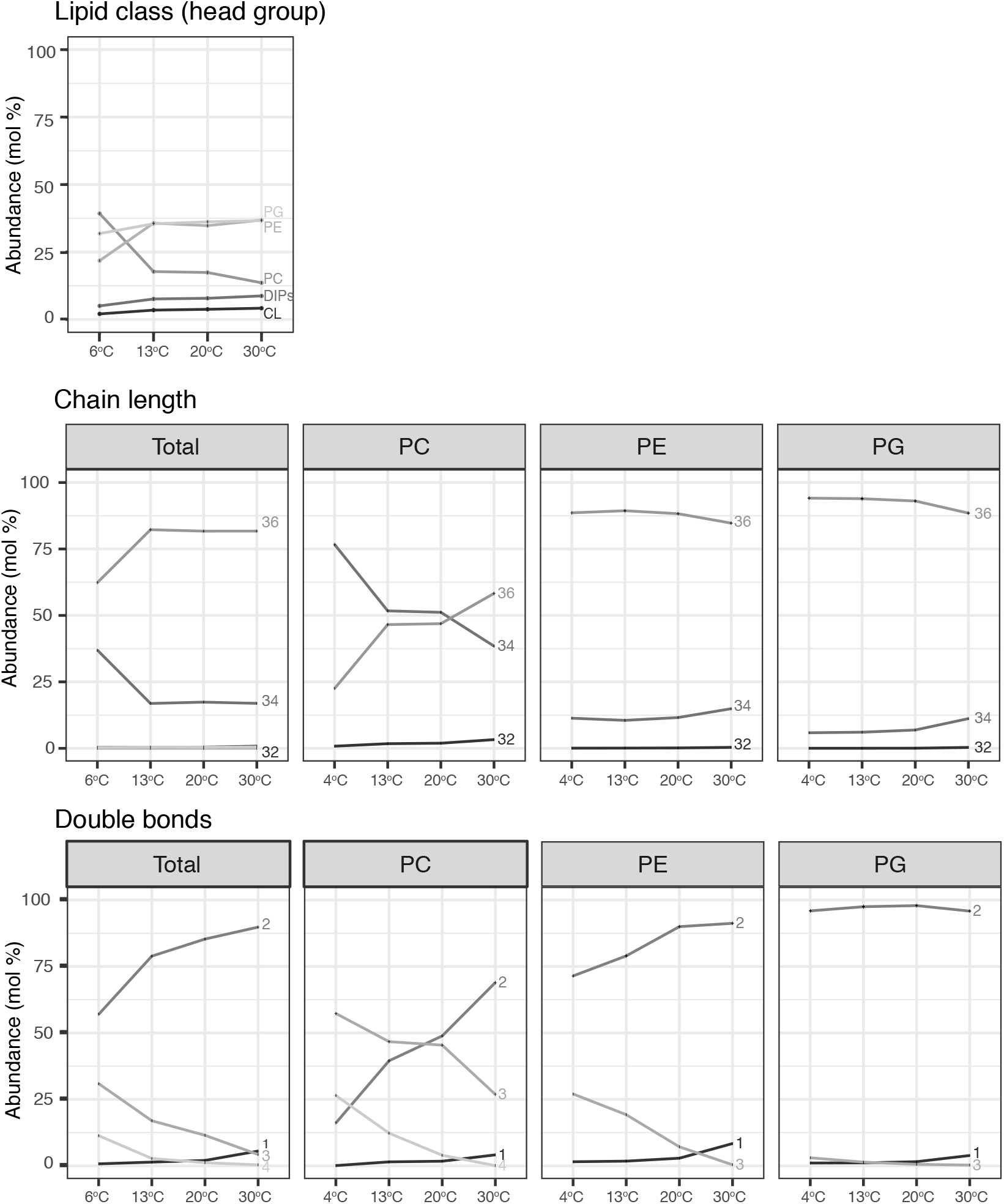
Step-wise adaptation of lipids by feature sampled at varying growth temperatures.

**Figure 6 – Supplement 3.**
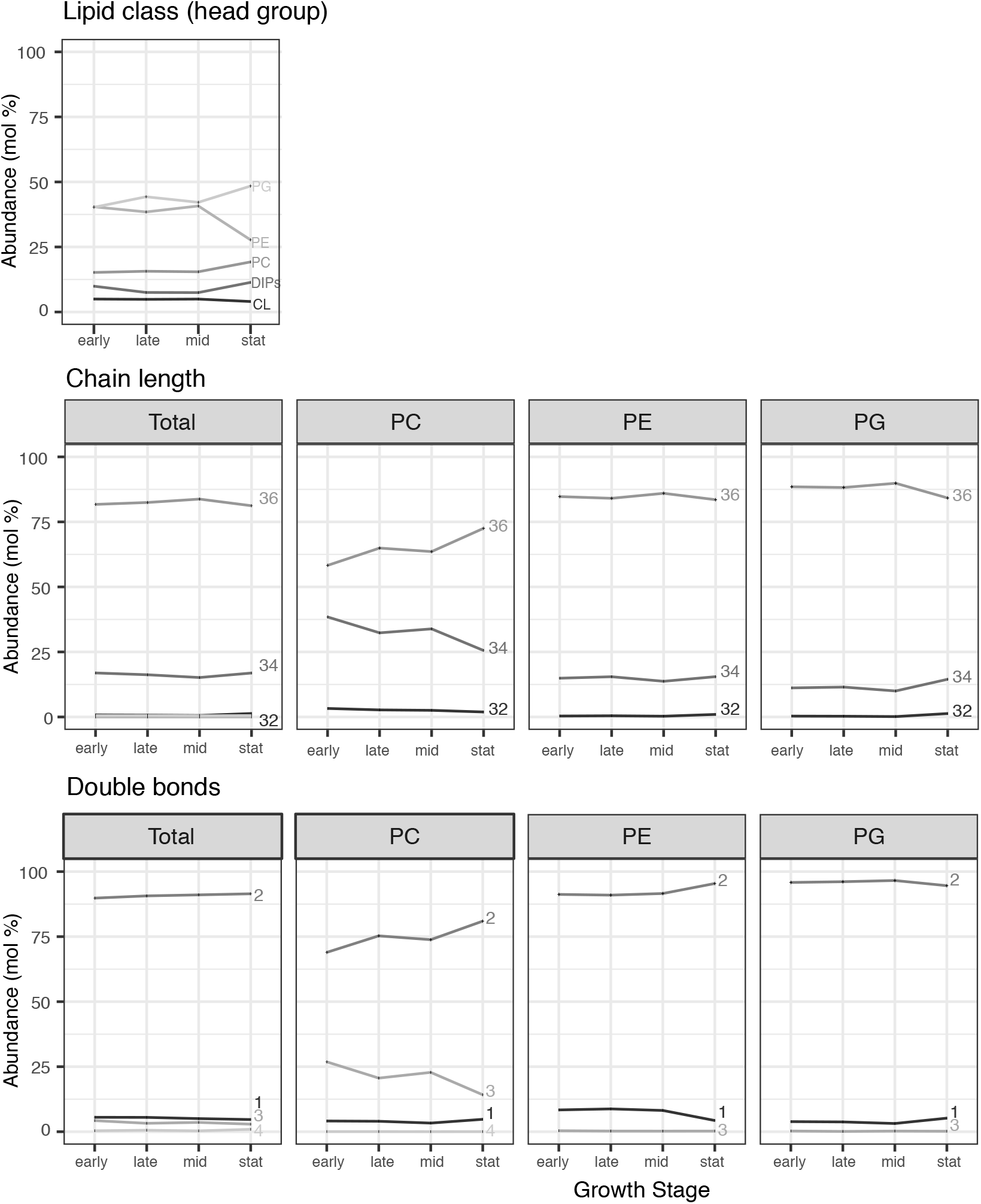
Step-wise adaptation of lipids by feature sampled at early, late, mid and stationary growth phase.

**Figure 6 – Supplement 4.**
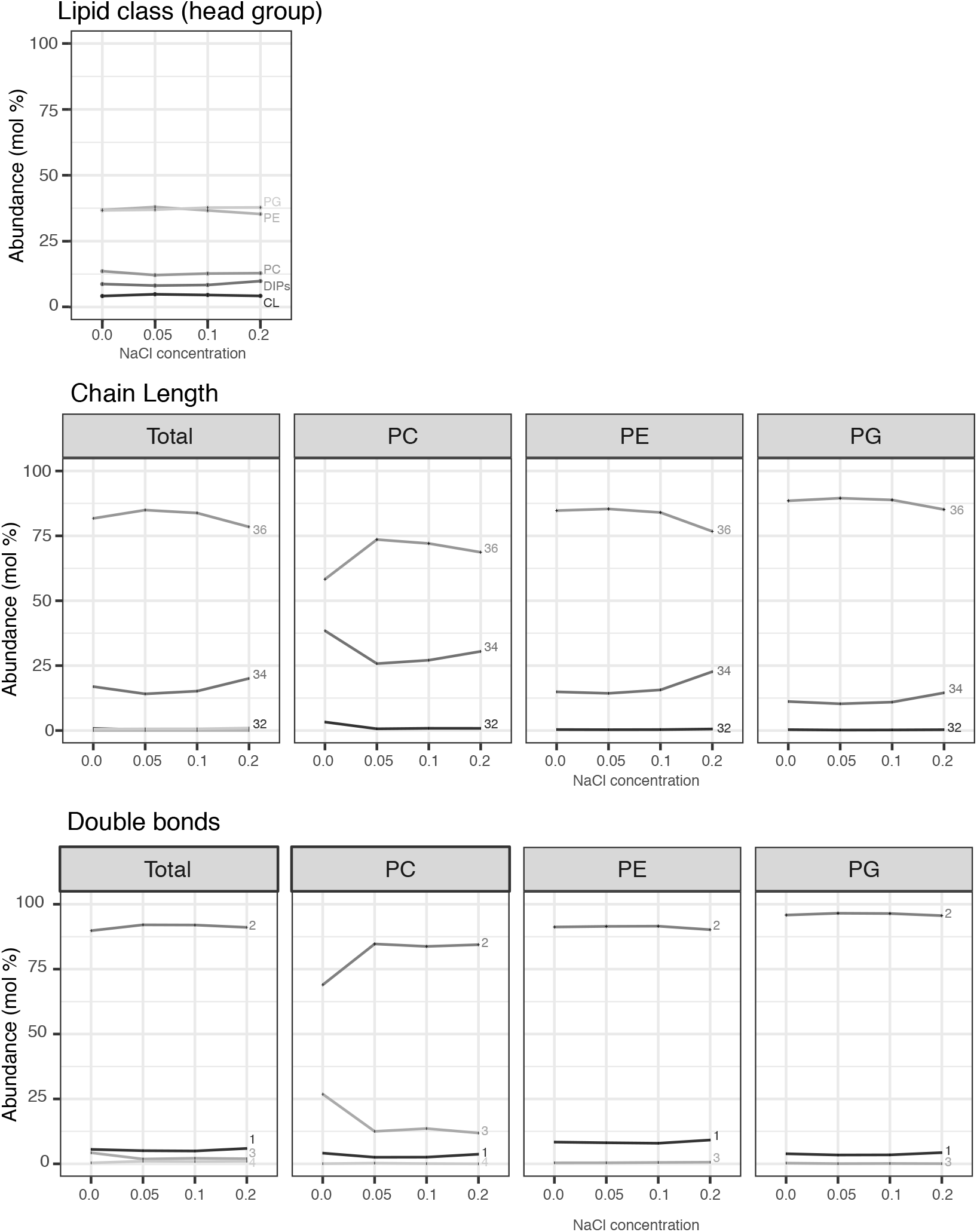
Step-wise adaptation of lipids by feature sampled at varying salt concentrations (NaCl).

